# Septins associate with AP-3 to support trafficking to the vacuole/lysosome in yeast

**DOI:** 10.64898/2026.02.13.705769

**Authors:** Mitchell Leih, Michaela McCright, Cortney Angers, Michael Davey, Elizabeth Conibear, Alex Merz, Greg Odorizzi

**Author notes:** Address correspondence to: Greg Odorizzi.

## Abstract

Adaptor protein complex 3 (AP-3) mediates clathrin-independent transport to lysosomes, yet accessory factors supporting this pathway remain incompletely defined. In *Saccharomyces cerevisiae*, the C-terminal intrinsically disordered regions (IDRs) of both AP-3 large subunits (δ and β3) serve as platforms for association with accessory factors. Through proteomic analysis of proteins associated with these IDRs, we identify the septin cytoskeleton as a candidate AP-3-associated factor. Bimolecular fluorescence complementation (BiFC) reveals a hierarchical pattern of association: AP-3 shows preferential proximity to core septin subunits (Cdc10, Cdc3, Cdc12) over terminal subunits (Cdc11 and Shs1). These terminal subunits serve as alternative caps of septin octamers, generating structurally distinct assemblies. Significantly, dysfunction of Cdc11 but not Shs1 selectively impairs AP-3-dependent cargo sorting without affecting the parallel vacuolar protein sorting (VPS) pathway to the vacuole (lysosome in yeast), providing genetic evidence for a specific functional connection between Cdc11-containing septin assemblies and AP-3-mediated transport.

## INTRODUCTION

Intracellular membrane trafficking relies on coat protein complexes that selectively package transmembrane protein cargoes into transport vesicles while recruiting auxiliary factors necessary for vesicle formation, targeting, and fusion with destination membranes (Bonifacino and Glick, 2004). Adaptor protein (AP) complexes are among the best-characterized vesicular coats. Of the five known AP complex family members, AP-1, AP-2, and AP-3 are most broadly conserved across eukaryotes (Hirst et al., 2011). While AP-1 and AP-2 function at the *trans*-Golgi network and plasma membrane respectively, AP-3 mediates transport from late Golgi/endosomal compartments to lysosomes and lysosome-related organelles (Cowles et al., 1997; Theos et al., 2005; Robinson, 2015). AP-3 is non-essential in most organisms, but its dysfunction in humans causes Hermansky-Pudlak syndrome, characterized by defects in pigmentation, hemostasis, and immune responses (Robinson, 2015; Dell’Angelica, 2009).

All AP complexes share a heterotetrameric architecture with two large subunits, a medium subunit, and a small subunit. The large subunits have C-terminal intrinsically disordered regions (IDRs) linked to N-terminal trunk regions, the latter of which form a compact core for membrane recognition and cargo selection (Kirchhausen et al., 2014). In AP-1 and AP-2, the IDRs terminate in ear domains that bind accessory proteins including clathrin (Edeling et al., 2006; Schmid et al., 2006). AP-3 exhibits distinct features, including a constitutively open conformation (Schoppe et al., 2021; Begley et al., 2024) and clathrin-independent function (Vowels and Payne, 1998; Zlatic et al., 2013; Schoppe et al., 2020). Moreover, ear domains are absent from IDRs in yeast AP-3, yet both IDRs are required for vesicle budding from late Golgi compartments (Leih et al., 2024), and at least one IDR directly binds the HOPS (homotypic fusion and protein sorting) complex that mediates tethering of AP-3 vesicles with vacuoles (yeast lysosomes) (Angers and Merz, 2009; Schoppe et al., 2020). These observations indicate that AP-3 IDRs serve as platforms for recruiting accessory proteins facilitating vesicular transport, yet the full repertoire of IDR-associating factors remains poorly defined.

In this study, we employed bimolecular fluorescence complementation (BiFC) and proteomic approaches to examine AP-3 interactions and spatial organization in yeast. We report that individual AP-3 complexes exhibit spatial proximity *in vivo*, and we identify the septin cytoskeleton as a functionally relevant AP-3 association in yeast. Our findings further reveal a hierarchical pattern of AP-3-septin interactions and demonstrate that specific septin components are required for AP-3-mediated cargo sorting, providing new insights into the organization of clathrin-independent trafficking to lysosomes/vacuoles.

## RESULTS AND DISCUSSION

### BiFC reveals spatial proximity between separate AP-3 complexes

To investigate AP-3 interactions *in vivo*, we employed bimolecular fluorescence complementation (BiFC), which reconstitutes Venus fluorescent protein when non-fluorescent N- and C-terminal fragments (VN and VC) are brought within ∼7 nm (Kerppola, 2006). VN or VC coding sequences were integrated in frame with the genomic *APL5* or *APL6* loci (encoding yeast δ and β3 large AP-3 subunits; Figure 1A) in haploid strains of opposite mating types. Mating these haploids generated heterozygous diploids coexpressing both VN and VC fusions alongside untagged wild-type alleles (Figure 1B). A chromogenic AP-3 cargo-sorting assay confirmed that all BiFC strains retained wild-type AP-3 function (Supplemental Figure S1).

**Figure 1.**
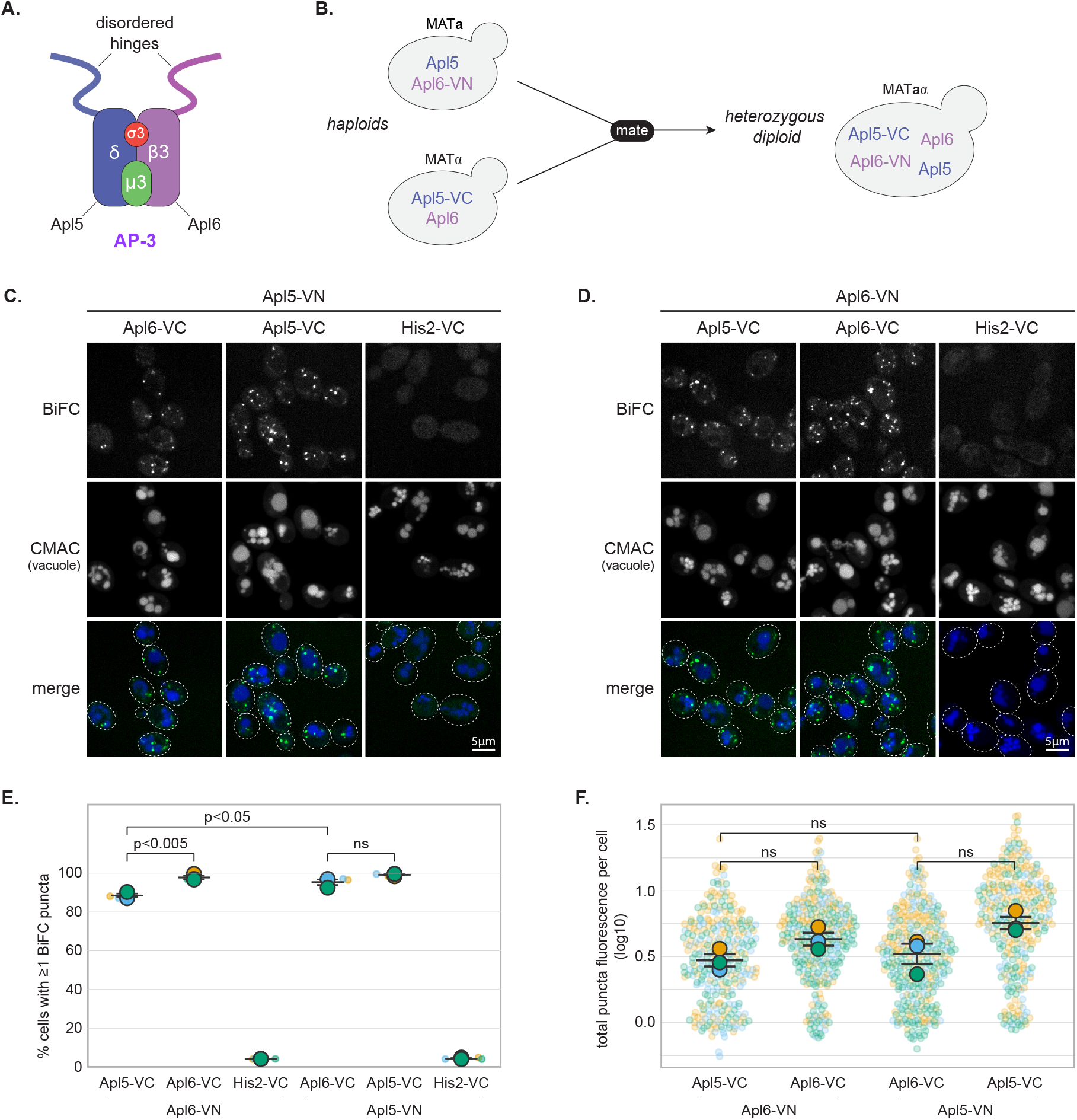
BiFC reveals spatial proximity between separate AP-3 complexes. **(A)** Schematic diagram of the AP-3 complex architecture. **(B)** Schematic diagram of the BiFC experimental design. VN or VC coding sequences were integrated in-frame at genomic *APL5* or *APL6* loci in haploid strains of opposite mating types. Also shown is the mating strategy to generate heterozygous diploids coexpressing BiFC fusions alongside wild-type alleles. **(C-D)** Representative confocal fluorescence microscopy images of BiFC strains showing heterotypic (Apl5-VN/Apl6-VC and reciprocal) and homotypic (Apl5-VN/Apl5-VC or Apl6-VN/Apl6-VC) interactions. His2-VC serves as negative control. Dashed ovals indicate cellular outlines. Scale bars, 5 μm. **(E)** Percentage of cells containing at least one BiFC punctum. Each small point represents the mean from one biological replicate; large symbols show means ± SEM. **(F)** Total puncta fluorescence per puncta-positive cell (log-transformed). Each small point represents one cell; large symbols show means ± SEM.

Robust puncta were observed by confocal microscopy in cells coexpressing Apl5-VN with Apl6-VC (Figure 1C) or Apl6-VN with Apl5-VC (Figure 1D), whereas His2-VC negative controls showed no fluorescence (Figure 1C-D). Surprisingly, “homotypic” pairings—Apl5-VN with Apl5-VC (Figure 1C) or Apl6-VN with Apl6-VC (Figure 1D)—produced similar BiFC puncta. Because individual AP-3 heterotetramers contain only one copy of each subunit, homotypic BiFC indicates that separate AP-3 complexes *in vivo* come into close proximity (within ∼7 nm). Quantitative analysis confirmed robust interactions (Figure 1E-F). Both heterotypic and homotypic pairings showed high frequencies of puncta-positive cells for both Apl5-based and Apl6-based combinations, with total puncta fluorescence per cell statistically indistinguishable between heterotypic and homotypic pairings.

BiFC detects proximity but does not distinguish between stable clustering versus repeated transient encounters at common membrane sites. Because BiFC is essentially irreversible once Venus reconstitution occurs (Finnigan et al., 2016; Garcia et al., 2016; Barve et al., 2018), signals accumulate over time, regardless of interaction stability. Thus, the BiFC signals derived from homotypic AP-3 subunit pairings likely reflect either stable spatial co-occupancy or dynamic recruitment to common membrane sites, potentially enhancing local AP-3 concentration for vesicle formation and tethering. We further validated the BiFC approach by confirming that it captures the established Vps41-Apl5 IDR interaction (Angers and Merz, 2009; Schoppe et al., 2020) and by demonstrating that the Apl5 IDR serves as the primary organizer of the AP-3-HOPS interface (Supplemental Figure S2).

### Proteomic identification of septins as AP-3 IDR-associated proteins

To identify novel AP-3-interacting proteins, we constructed recombinant GST fusions to the Apl5 IDR (residues 711-932) or Apl6 IDR (residues 744-809), expressed them in bacteria, and incubated them with yeast lysates (Figure 2A). Mass spectrometry of bound proteins identified all five subunits of the mitotic septin complex (Figure 2B and Supplemental Table S1). Septins are conserved GTPases forming hetero-oligomeric filaments essential for membrane organization (Pan et al., 2007; Bertin et al., 2008; Caudron and Barral, 2009). The five mitotic septin subunits in yeast (Cdc3, Cdc10, Cdc11, Cdc12, Shs1) organize the cytokinetic contractile ring and actin cytoskeleton (Hartwell, 1971; Mino et al., 1998; Bi et al., 1998; McMurray et al., 2011; Longtine et al., 2000), though septins in diverse organisms also function in membrane trafficking and other processes (Spiliotis and Gladfelter, 2012). Septin subunits were identified with substantial sequence coverage (3.6-15.8%; Figure 2B). Apl5 IDR recovered Cdc10, Cdc3, Cdc11, and Shs1; Apl6 IDR recovered Cdc3 only; combined IDRs recovered Cdc10, Cdc3, and Cdc12.

**Figure 2.**
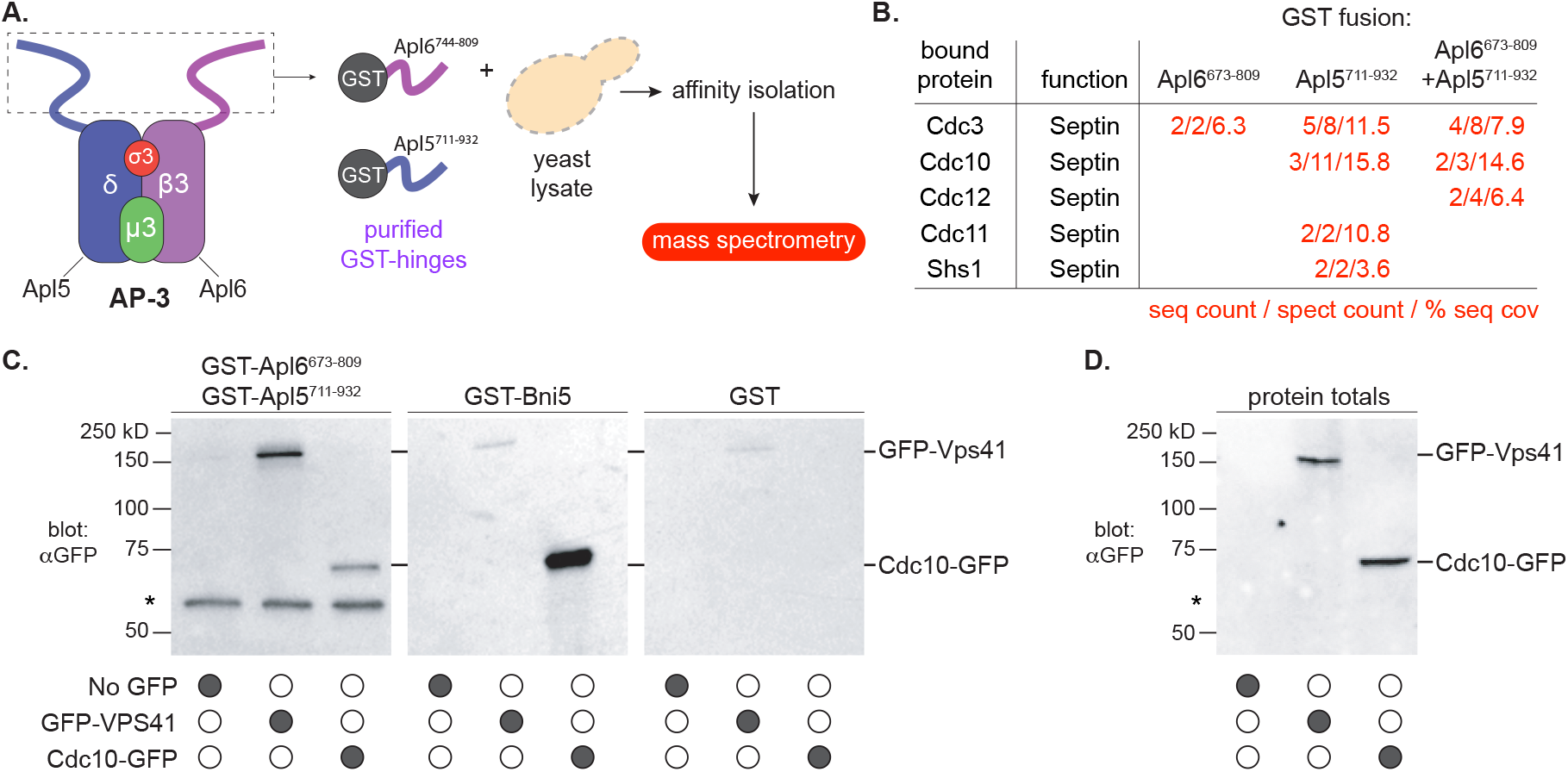
Proteomic identification of septins as AP-3 IDR-associated proteins. **(A)** Schematic of GST pulldown experimental design. GST fusions to Apl5 IDR (residues 711-932) or Apl6 IDR (residues 744-809) were expressed in bacteria, purified on glutathione resin, and incubated with yeast whole-cell lysates. After washing, copurifying proteins were identified by mass spectrometry. **(B)** Mass spectrometry results showing sequence count, spectrum count, and percent sequence coverage for the five mitotic septin subunits (Cdc3, Cdc10, Cdc11, Cdc12, Shs1) recovered from GST-IDR pulldowns. Complete proteomic data in Supplemental Table S1. **(C)** Western blot validation of septin-AP-3 IDR interactions. GST alone (negative control), GST-IDRs (Apl5+Apl6 combined), or GST-Bni5 (positive control for septin binding) were incubated with lysates from cells expressing GFP alone, Cdc10-GFP, or GFP-Vps41 (positive control for AP-3 IDR binding), then analyzed by anti-GFP immunoblotting. Input lanes show 5% of lysate used in pulldowns. **(D)** Western blot analysis of protein amounts in total fractions used for pulldowns shown in (C).

Western blot validated the findings. Lysates expressing GFP alone, Cdc10-GFP, or GFP-Vps41 were incubated with GST alone, GST-IDRs (Apl5 and Apl6 combined), or GST-Bni5 (a known septin-interacting protein; Lee et al., 2002). AP-3 IDRs pulled down Cdc10-GFP and GFP-Vps41; GST alone pulled down neither (Figure 2C-D). Because septin subunits form obligate hetero-oligomeric complexes (Field et al., 1996; Frazier et al., 1998; Bertin et al., 2008), Cdc10-GFP detection likely represents copurification of intact septin octamers.

Mammalian septins SEPT6 and SEPT7 (orthologs of yeast Cdc12 and Cdc3) were previously recovered in AP-3 proteomic screens (Baust et al., 2008; Traikov et al., 2014), suggesting conserved septin-AP-3 association. Our analysis (Supplemental Table S1) also identified septin regulatory factors (Hsl7; Shulewitz et al., 1999), the septin-associated kinase Cla4 (Versele and Thorner, 2004), and actin cytoskeletal components (Abp1, Abp140, Act1, Tpm1, Rvs167). The recovery of these established septin-associated proteins validates that intact septin complexes were captured by AP-3 IDRs (Hartwell, 1971; Longtine et al., 2000). Because our pulldowns were performed from whole-cell lysates, we cannot determine whether septins bind AP-3 IDRs directly or indirectly. However, as shown below, genetic analysis demonstrates that specific septin components are functionally required for AP-3-mediated trafficking, providing validation beyond biochemical association. The identification of all five septin subunits along with established septin-interacting proteins indicates that AP-3 IDRs associate with assembled septin complexes.

### BiFC reveals hierarchical AP-3 interactions with septin proteins

The five mitotic septin proteins in yeast assemble into palindromic octamers (Figure 3A; Bertin et al., 2008; Garcia et al., 2011) with central Cdc10 homodimers flanked by Cdc3 and Cdc12, and either Cdc11 or Shs1 occupying terminal positions (Bertin et al., 2008; Garcia et al., 2011; McMurray and Thorner, 2019). Cdc11-based octamers polymerize into straight filaments, whereas Shs1-based octamers form curved bundles (Garcia et al., 2011; Booth et al., 2015). We tested AP-3 proximity to each septin using BiFC.

**Figure 3.**
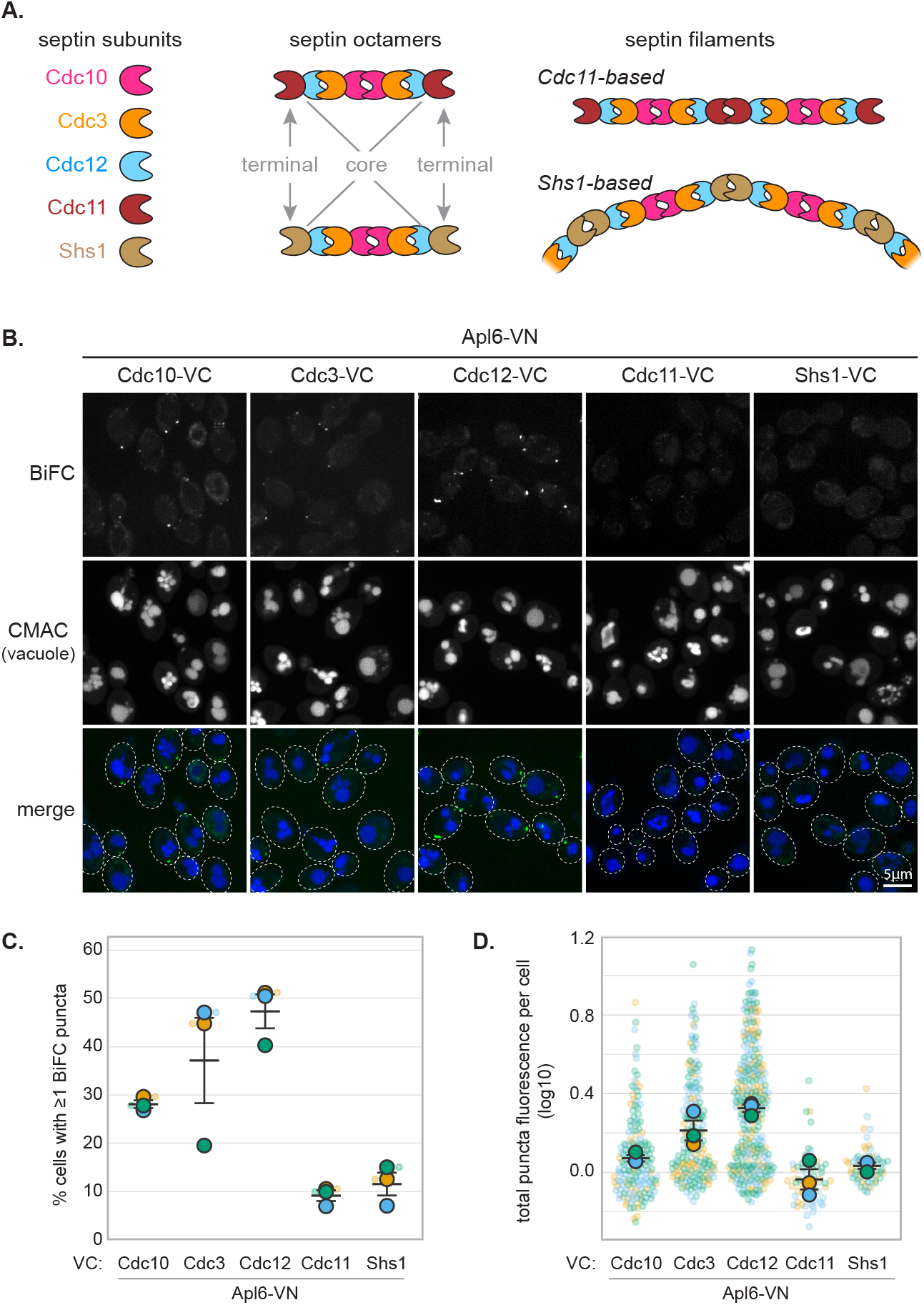
BiFC reveals hierarchical AP-3 proximity to septin proteins. **(A)** Schematic diagram of septin octamer organization showing palindromic arrangement. Core subunits (Cdc10, Cdc3, Cdc12) form the central scaffold, while terminal subunits (Cdc11 or Shs1) occupy end positions and determine higher-order filament assembly properties. Cdc11-based octamers polymerize end-to-end into straight paired filaments; Shs1-based octamers form curved, laterally associated bundles and rings. **(B)** Representative confocal microscopy images showing BiFC between Apl6-VN and indicated septin-VC fusions. Note robust puncta with core septins (Cdc10, Cdc3, Cdc12) and diminished signal with terminal septins (Cdc11, Shs1). Dashed ovals indicate cellular outlines. Scale bars, 5 μm. **(C)** Percentage of cells containing at least one BiFC punctum. **(D)** Total puncta fluorescence per puncta-positive cell. Statistical comparisons are reported in the main text and omitted from the graphs in (C) and (D) to avoid clutter.

Septin-VC fusions were coexpressed in diploid cells with Apl6-VN or Apl5-VN versus His2-VN. Core septin fusions (Cdc10-VC, Cdc3-VC, Cdc12-VC) paired with Apl6-VN (Figure 3B) or Apl5-VN (Supplemental Figure S3A) produced fluorescent puncta, with Cdc12-VC showing the highest frequency of puncta-positive cells, followed by Cdc10-VC, then Cdc3-VC (Figure 3C). No septin-VC fusion produced significant fluorescent puncta with His2-VN (Supplemental Figure S3B). Notably, the alternative terminal octamer subunits (Cdc11 and Shs1) showed markedly reduced BiFC signals when paired with AP-3, with puncta frequencies barely distinguishable from His2-VN negative controls (Cdc11-VC: p = 0.064; Shs1-VC: p = 0.102) and significantly lower than the core septin Cdc12-VC (p = 0.005 and p = 0.002, respectively; Figure 3C). Similarly, both terminal septins showed reduced puncta fluorescence compared to Cdc12-VC (p = 0.012 and p = 0.0003; Figure 3D). Both Cdc11-VC and Shs1-VC produced robust BiFC fluorescence when coexpressed with their neighboring septin Cdc12-VN (Weems and McMurray, 2017; our unpublished observations), demonstrating these fusion proteins are properly expressed and assembled into septin octamers.

BiFC signal intensity reflects fluorophore reconstitution efficiency within constrained protein assemblies and need not correlate with functional requirement. The reduced BiFC signal with terminal septins therefore reflects genuine differences in proximity or accessibility rather than indicating these subunits are functionally unimportant. The hierarchy across both AP-3 subunits and all septin positions supports a model in which AP-3 engages assembled septin octamers with preferential access to core elements.

### Specific septin mutations selectively impair AP-3-dependent cargo sorting

To investigate the functional relevance septins have to AP-3 trafficking, we examined the transport of GNSS, an AP-3 reporter cargo protein we previously characterized (Plemel et al., 2021; Leih et al., 2024). The essential septin genes *CDC10, CDC3, CDC12*, and *CDC11* (Hartwell, 1971) were analyzed using point-mutant alleles; the non-essential *SHS1* gene was deleted (Carroll et al., 1998; Mino et al., 1998; Garcia et al., 2011). A chromogenic GNSS sorting assay based on secreted invertase activity (Darsow et al., 2000; Burston et al., 2009; Dalton et al., 2015) showed that *cdc10-5, cdc12-td* (temperature-sensitive degron; Li et al., 2011; Weems et al., 2014), and *cdc11-4* caused pronounced GNSS mislocalization comparable to *apl6*Δ (Figure 4A). Weaker mislocalization resulted from *cdc10-4, cdc11-1*, and *cdc11-3*. GNSS localization remained normal in *shs1*Δ cells, demonstrating AP-3 transport specifically depends on Cdc11-capped octamers.

**Figure 4.**
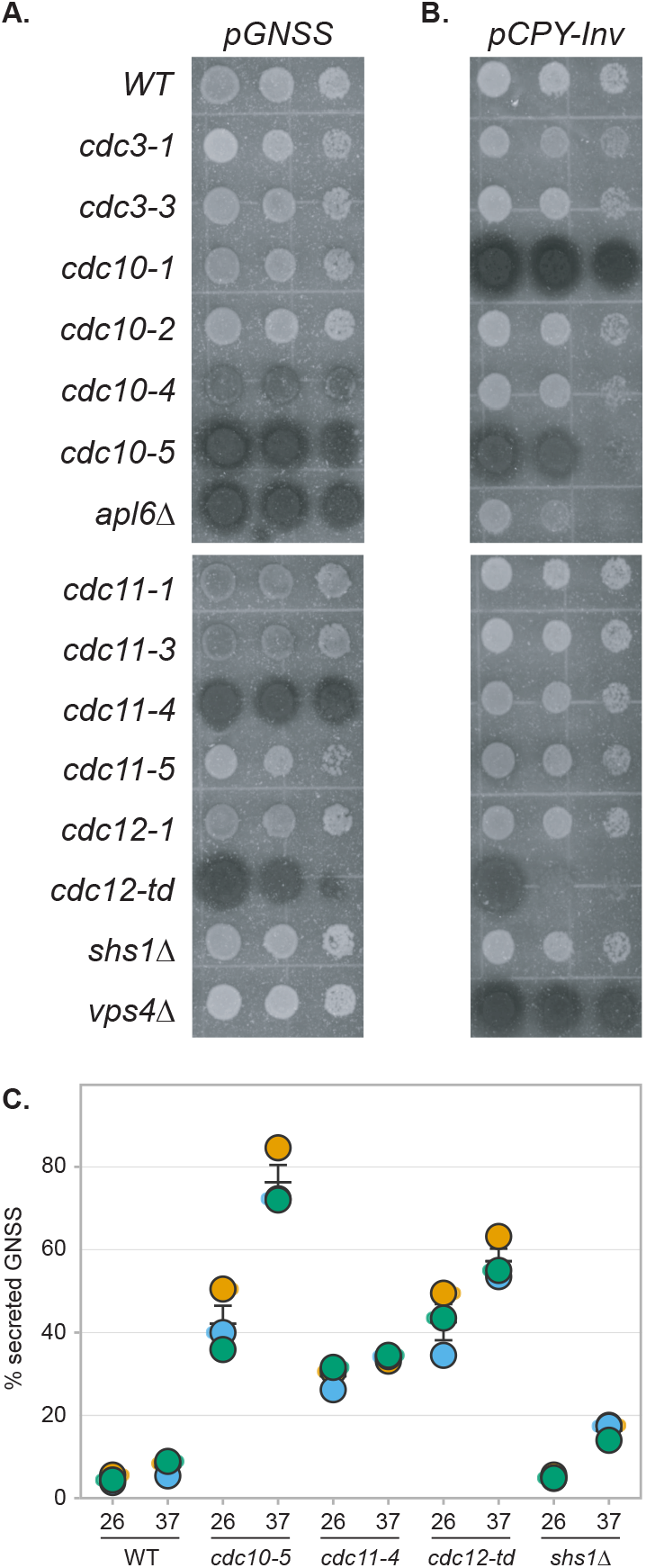
Specific septin mutations selectively impair AP-3-dependent cargo sorting. **(A)** Chromogenic overlay assay showing GNSS (AP-3 cargo) localization in indicated septin mutant strains. Darker coloration indicates greater cell-surface invertase activity due to GNSS mislocalization to the plasma membrane. *apl6*Δ serves as positive control for AP-3 pathway defect. (B) Chromogenic overlay assay showing CPY-Invertase (VPS pathway cargo) localization in the same strains, demonstrating pathway selectivity. The *cdc11-4* allele does not affect CPY-Invertase sorting (comparable to WT), whereas *cdc10-5, cdc12-td*, and *cdc10-1* cause CPY-Invertase mislocalization. *vps4*Δ serves as positive control for VPS pathway defect. **(C)** Quantitative liquid invertase assay measuring percentage of secreted GNSS invertase activity. Cells were grown to mid-logarithmic phase at 26°C, then maintained at 26°C or shifted to 37°C for 1 hour before harvesting. Secretion calculated as (supernatant activity / total activity) × 100%. Bars show means ± SEM from n=3 independent biological replicates. Statistical comparisons were performed using one-way ANOVA; p-values are reported in the main text for comparisons to wild-type at the same temperature and omitted from the graph to avoid clutter.

We also examined the trafficking of CPY-Invertase, a reporter cargo protein of the VPS pathway to the vacuole (Bankaitis et al., 1986), which operates in parallel with the AP-3 pathway and is genetically distinct (Cowles et al., 1997). Because septins function in multiple processes (Hartwell, 1971; Barral et al., 2000; Caudron and Barral, 2009), general dysfunction should affect both CPY-Invertase and GNSS, whereas AP-3-specific defects should affect only GNSS. The *cdc10-5* and *cdc12-td* mutations caused pronounced CPY-Invertase mislocalization (Figure 4B), indicating broad vacuolar trafficking defects. In striking contrast, *cdc11-4* did not affect CPY-Invertase localization (Figure 4B), indicating selective AP-3 impairment. Similarly, *shs1*Δ exhibited normal CPY-Invertase sorting (Figure 4B).

Quantitative liquid invertase assays confirmed these defects (Figure 4C). At 26°C, wild-type cells secreted 5±1% of total GNSS. In contrast, *cdc10-5* and *cdc12-td* cells secreted 42±4% (approximately 8-fold increase, p=0.012), and *cdc11-4* cells secreted 29±2% (approximately 6-fold increase, p=0.002). Temperature shift to 37°C significantly enhanced sorting defects in *cdc10-5* (76±4%, p=0.002 vs. wild-type at 37°C) and *cdc12-td* (57±3%, p=0.001). Notably, *shs1*Δ cells showed no significant GNSS mislocalization at 26°C (5±0%, p=0.46), though a minor increase was observed at 37°C (16±1%, p=0.005).

The selective AP-3 trafficking defect in *cdc11-4* cells without VPS impairment (Figure 4B) supports specific Cdc11 function in AP-3 transport, whereas the broad trafficking defects in *cdc10-5* and *cdc12-td* strains likely reflect general septin destabilization (Weems et al., 2014; Barve et al., 2018). Cdc11 exhibits weak BiFC proximity to AP-3 yet is specifically required for AP-3 sorting, demonstrating BiFC signal strength does not correlate with functional importance for architectural components. As with regulatory subunits in cohesin and other structural complexes that organize architecture without directly contacting substrates (Haering et al., 2002), Cdc11 occupies the terminal octamer position and is critical for end-to-end polymerization (Bertin et al., 2008; McMurray et al., 2011), organizing septin filament architecture that supports AP-3 function without requiring direct AP-3 contact. The *cdc11-4* allele contains two amino acid substitutions (S31F and S100P) that alter the P-loop critical for nucleotide binding (Weems et al., 2014), likely perturbing structural stability and disrupting higher-order septin assembly. The allele-specific separation of AP-3 and VPS pathway phenotypes indicates that proper septin filament architecture, rather than general septin function, is specifically required for AP-3-mediated transport.

We propose a working model in which septins could concentrate AP-3 within specific membrane domains, create barriers regulating vesicle motility, or organize territories facilitating tethering/fusion. These possibilities are not mutually exclusive and our data do not distinguish between them. However, all three models share a common framework in which septins organize higher-order membrane architecture that spatially regulates AP-3 function. This framework is supported by three observations: AP-3 shows preferential proximity to core septin subunits forming the architectural scaffold; the *cdc11-4* allele, which disrupts higher-order septin filament assembly, selectively impairs AP-3 but not VPS trafficking; and septins are established organizers of membrane domains (Spiliotis and Gladfelter, 2012; Bridges and Gladfelter, 2015). Future work employing localization studies and analysis of AP-3 vesicle dynamics in septin mutants will be required to distinguish between these and other possible mechanisms.

## MATERIALS AND METHODS

### Construction of yeast strains and DNA plasmids

Standard techniques were used for growth and genetic manipulation of *S. cerevisiae* strains and construction of plasmids (Supplemental Table S1). Yeast strains were constructed by one-step PCR-based integration (Longtine et al., 1998). Diploid strains were generated by mating haploid strains of opposite mating types and selecting on appropriate media. Strains were verified by PCR analysis of genomic DNA.

Bacterial expression plasmids encoding GST fused to Apl5 IDR (residues 711-932) or Apl6 IDR (residues 744-809) were constructed by PCR amplification, digestion with NcoI and BamHI, and ligation into NcoI/BamHI-digested pGST-Parallel (Sheffield et al., 1999). Plasmid pTEF1-VN-416 was constructed by homologous recombination of SacI-digested pRS416 (Sikorski and Hieter, 1989) with overlapping DNA fragments encoding the TEF1 promoter and VN coding sequence. Plasmid pTEF1-VN-VPS41 was constructed by homologous recombination of AscI-digested pTEF1-VN-416 with an overlapping DNA fragment encoding *VPS41*. Plasmids were verified by DNA sequencing. Yeast and DNA reagents are available upon request.

### Strains and plasmids used for BiFC and trafficking assays

BiFC analysis of AP-3 subunit proximity (Figure 1) was performed using strains GOY1137, GOY1138, GOY1168, GOY1188, GOY1190, and GOY1193. BiFC analysis of AP-3-septin proximity (Figure 3) was performed using strains GOY1199, GOY1200, GOY1201, GOY1220, and GOY1227. Cargo sorting assays (Figure 4) were performed using control and septin-mutant strains with the indicated genotypes. All strains used for GNSS trafficking assays contain *suc2*Δ to eliminate endogenous invertase expression and were transformed with plasmid pLC1514 encoding the GNSS reporter cargo protein (Leih et al., 2024).

### Fluorescence microscopy and BiFC quantification

Liquid cultures were grown at 30°C to logarithmic phase before staining endosomal membranes with 1.6 μM FM4-64 (Invitrogen) for 25 min followed by a 90 min chase in stain-free YPD (Odorizzi et al., 2003; Vida and Emr, 1995). Live cells were observed at room temperature with an inverted fluorescence microscope (Ti2-E PSF; Nikon) equipped with a Yokogawa CSU-X1 spinning disk confocal system and a 100× numerical aperture 1.45 oil objective (Plan Apo λ; Nikon). Images were acquired using an Andor iXon Ultra 512×512 EMCCD camera with Micromanager version 2.0 software and analyzed with ImageJ (NIH).

For BiFC quantification, maximum-intensity projections of confocal z-stacks were acquired under identical settings for all strains within each experiment. BiFC fluorescence was quantified using Fiji (ImageJ; NIH) with a custom macro. Maximum-intensity projections of confocal z-stacks acquired under identical settings were analyzed. To establish detection thresholds, negative-control strains expressing His2-VN paired with the corresponding VC-tagged protein were imaged in parallel. Using Fiji’s Find Maxima function, a threshold was set for each imaging session to exclude the majority of visually apparent puncta in His2 controls while retaining experimental puncta. Individual cells were manually segmented using region-of-interest (ROI) outlines, excluding cells with abnormally high diffuse autofluorescence indicative of compromised membrane integrity. Puncta were assigned to cells based on spatial containment within ROIs; puncta outside live-cell ROIs were excluded. For each punctum, integrated fluorescence intensity was measured within a fixed-radius circular region centered on the punctum coordinate. Total puncta fluorescence per cell was calculated by summing integrated intensities of all puncta within that cell. To normalize for day-to-day variation, total puncta fluorescence per cell was divided by the median total puncta fluorescence measured in the His2 control from the same imaging session. Normalized data from independent imaging sessions were visualized using SuperPlotsOfData—a web app for transparent display and quantitative comparison of continuous data (Goedhart, 2021). For each experimental condition, cells were scored for puncta presence per biological replicate, and puncta fluorescence was quantified from puncta-positive cells per replicate across n=3 independent biological replicates. Statistical comparisons were performed using one-way ANOVA with Tukey’s post-hoc test. For all BiFC experiments, n=3 independent biological replicates were analyzed, with ≥100 cells scored per replicate for puncta presence (≥50 cells per replicate in Supplemental Figure S2).

### GST pulldown assays, western blotting, and mass spectrometry

*E. coli* BL21(DE3) cells transformed with pGST-Parallel vectors encoding GST fusions to Apl5 IDR (residues 711-932) or Apl6 IDR (residues 744-809) were grown to OD□ □ □ ∼1.5 in Terrific Broth at 37°C. Protein expression was induced with 200 μM IPTG. Cells expressing GST-Apl6-IDR were incubated overnight at 18°C; cells expressing GST or GST-Apl5-IDR were incubated 3 hours at 37°C. Cells were pelleted, resuspended in 40 mL bacterial lysis buffer (PBS pH 7.4, 1 mM DTT, 1 mM PMSF, 1 μg/mL leupeptin, 1 μg/mL pepstatin), dripped into liquid nitrogen, and stored at −80°C.

To prepare GST-affinity resins, frozen cell pellets (∼700 μL packed volume after thaw) were supplemented with 0.5% Triton X-100 and 10 μg DNase I, lysed by sonication, and centrifuged at 20,000 × *g* for 20 min to produce clarified lysate. Five hundred microliters of lysate were added to 50 μL Glutathione Sepharose 4B (Cytiva, Marlborough, MA) pre-washed with PBS and incubated overnight at 4°C with rotation. Beads were washed three times each with 500 μL PBS, 500 μL PBS containing 350 mM NaCl, and 500 μL yeast lysis buffer (20 mM HEPES pH 6.8, 0.2 M sorbitol, 2 mM EDTA, 50 mM potassium acetate, 1 μg/mL aprotinin, 1 μg/mL leupeptin, 1 μg/mL pepstatin, 1 μg/mL Pefabloc-SC, 1 mM PMSF).

Yeast cell lysates (wild-type strain BY4742; Brachmann et al., 1998) were prepared from 1-liter cultures grown to mid-logarithmic phase. Cells were converted to spheroplasts, gently pelleted at 1,000 × *g* for 2 min, resuspended in yeast lysis buffer, and homogenized by 25 strokes in a Dounce homogenizer. Lysates were clarified by centrifugation at 1,000 × *g* for 5 min, supplemented with 0.5% Triton X-100, and centrifuged at 20,000 × *g* for 15 min to remove insoluble material. Washed GST resin was incubated with 500 μL (∼150 OD□ □ □ units) of yeast detergent lysate for 1 hour at 4°C with rotation, then washed three times with yeast lysis buffer.

For western blot analysis, proteins were eluted with glutathione elution buffer (50 mM Tris-Cl pH 7.9, 20 mM reduced glutathione, 600 mM NaCl, 1% Triton X-100), resolved by SDS-PAGE, transferred to nitrocellulose, and immunoblotted with anti-GFP antibodies (Roche, Basel, Switzerland).

For mass spectrometry, GST pulldowns were performed as above with modifications. Triton X-100 was omitted from final washes. Proteins were eluted with 100 μL elution buffer (50 mM ammonium bicarbonate pH 7.8, 0.5 M NaCl, 0.1% RapiGest SF; Waters Corporation, Milford, MA) and stored at −80°C. Thawed samples were diluted 1:1 in 50 mM ammonium bicarbonate pH 7.8 containing 0.1% RapiGest, boiled at 99°C for 2 min, reduced with 5 mM DTT at 60°C for 30 min, and alkylated with 15 mM iodoacetamide in the dark at room temperature for 30 min. Samples were digested overnight with trypsin (Promega, Madison, WI) at 37°C, acidified with HCl to 100 mM final concentration, and incubated at 37°C for 45 min. Samples were centrifuged at 13,000 × *g* for 10 min at 4°C, and supernatants were analyzed by LC-MS/MS using an LTQ mass spectrometer (Thermo Fisher Scientific, Waltham, MA). Results were processed with SEQUEST and DTASelect software. Complete mass spectrometry data including all identified proteins, sequence coverage values, and functional annotations are provided in Supplemental Table S1.

### Invertase activity assays

Chromogenic overlay assays for cargo sorting were performed as described (Darsow et al., 2000; Burston et al., 2009; Dalton et al., 2015). Cells were spotted to agar medium in which fructose was substituted for glucose as the carbon source and incubated at 26°C for 2 days. Plates were then overlaid with top agar containing chromogenic solution (125 mM ultrapure sucrose, 166 mM sodium acetate pH 5.2, 0.666 mM N-ethylmaleimide, 0.017 mg/mL horseradish peroxidase, 15 units/mL glucose oxidase, 1 mg/mL o-dianisidine, 3% w/v agar). Secreted invertase activity appears as a color change from white to reddish-brown within ∼15 min, indicating mislocalization of cargo to the cell surface. To verify equivalent cargo expression across strains, parallel overlay assays were performed on cells exposed to chloroform vapor to permeabilize membranes and release intracellular invertase. Images shown in Figure 4A-B are representative of n=3 independent experiments.

For quantitation of secreted invertase activity, cultures were grown to mid-logarithmic phase at 26°C, then maintained at 26°C or shifted to 37°C for 1 hour. Cells (0.4 OD□ □ □ units) were centrifuged, washed twice in 100 mM sodium acetate pH 5.2, and resuspended in 400 μL of 100 mM sodium acetate pH 5.2. Samples were split into two groups of 190 μL each to measure extracellular and total invertase activity using the liquid invertase assay (Darsow et al., 2000). Measurements were performed in triplicate for each of three independent biological replicates. Statistical comparisons were performed using one-way ANOVA with SuperPlotsOfData (Goedhart, 2021).

## ACKNOWLEDGEMENTS

We thank Erfei Bi (University of Pennsylvania) for providing the plasmid encoding GST-Bni5.

## FIGURE LEGENDS

**Supplemental Figure S1.**
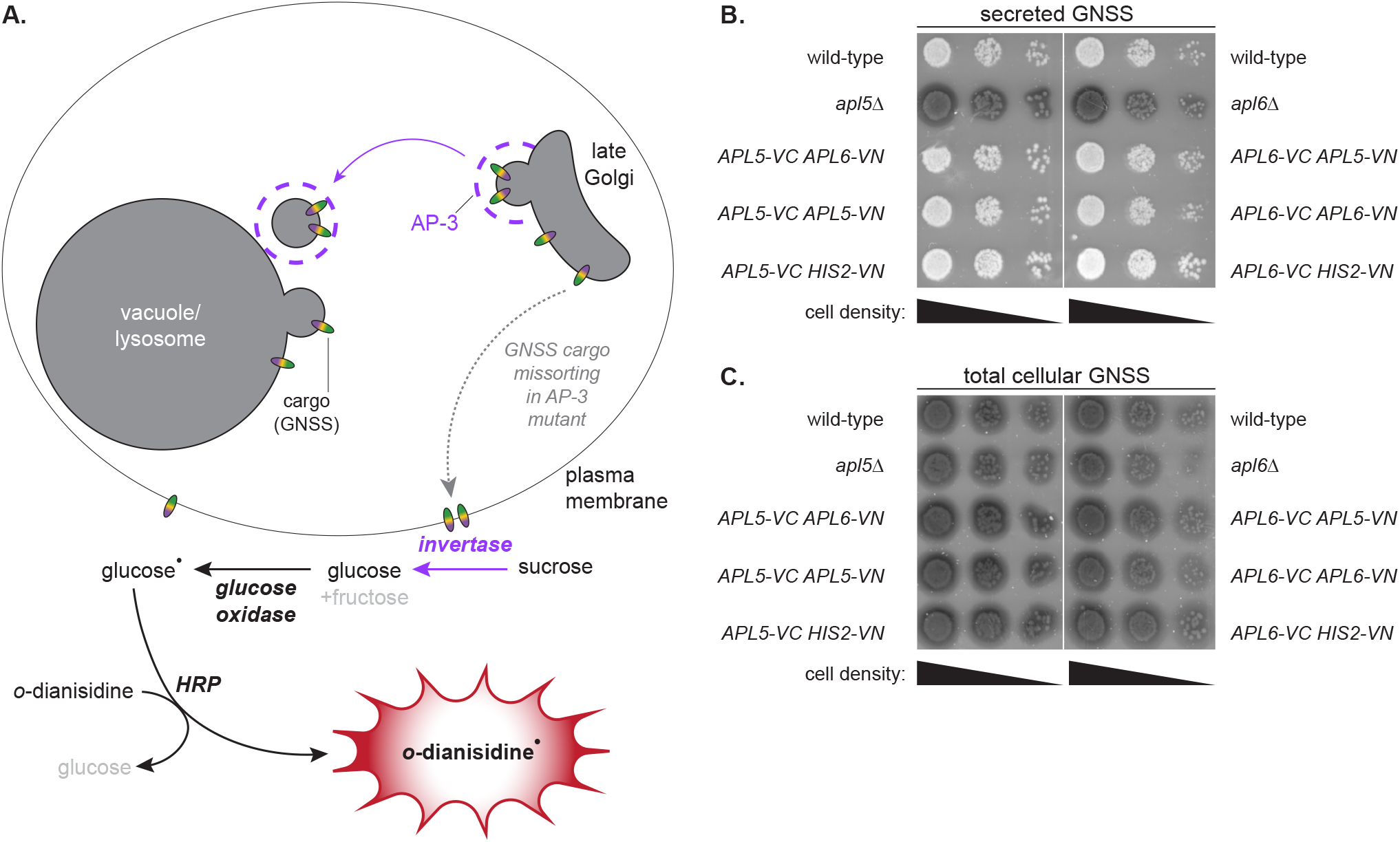
BiFC fusions preserve AP-3 function. To assess whether BiFC fusions compromised AP-3 function, we examined trafficking of GNSS, a synthetic AP-3 cargo protein reporter that contains two invertase domains fused to its exoplasmic domain (Plemel et al., 2021; Leih et al., 2024). (A) Schematic diagram explaining the chromogenic invertase activity assay principle. GNSS normally traffics from the Golgi to the vacuole membrane via the AP-3 pathway. When AP-3 function is compromised, GNSS mislocalizes to the plasma membrane, where cell-surface invertase activity can be detected chromogenically using an overlay assay (Darsow et al., 2000). (B) Chromogenic overlay assay results for all BiFC strains used in Figure 1, showing normal GNSS internalization (lack of dark coloration) comparable to wild-type control; *apl5Δ* and *apl6Δ* serve as positive controls showing GNSS mislocalization (dark coloration). Representative images from n=3 independent experiments demonstrate that BiFC fusions—including both heterotypic pairings (Apl5-VN/Apl6-VC and reciprocal Apl6-VN/Apl5-VC) and homotypic pairings (Apl5-VN/Apl5-VC and Apl6-VN/Apl6-VC)—do not impair AP-3-dependent cargo sorting. The preservation of normal GNSS trafficking in cells with homotypic BiFC combinations demonstrates that irreversible Venus reconstitution does not trap AP-3 complexes in non-functional aggregates; rather, AP-3 retains the dynamic properties necessary for productive cargo sorting even after BiFC capture. (C) Chromogenic assay of the same cells shown in (B) after cellular lysis upon exposure to chloroform vapor, confirming equivalent GNSS expression across all strains.

**Supplemental Figure S2.**
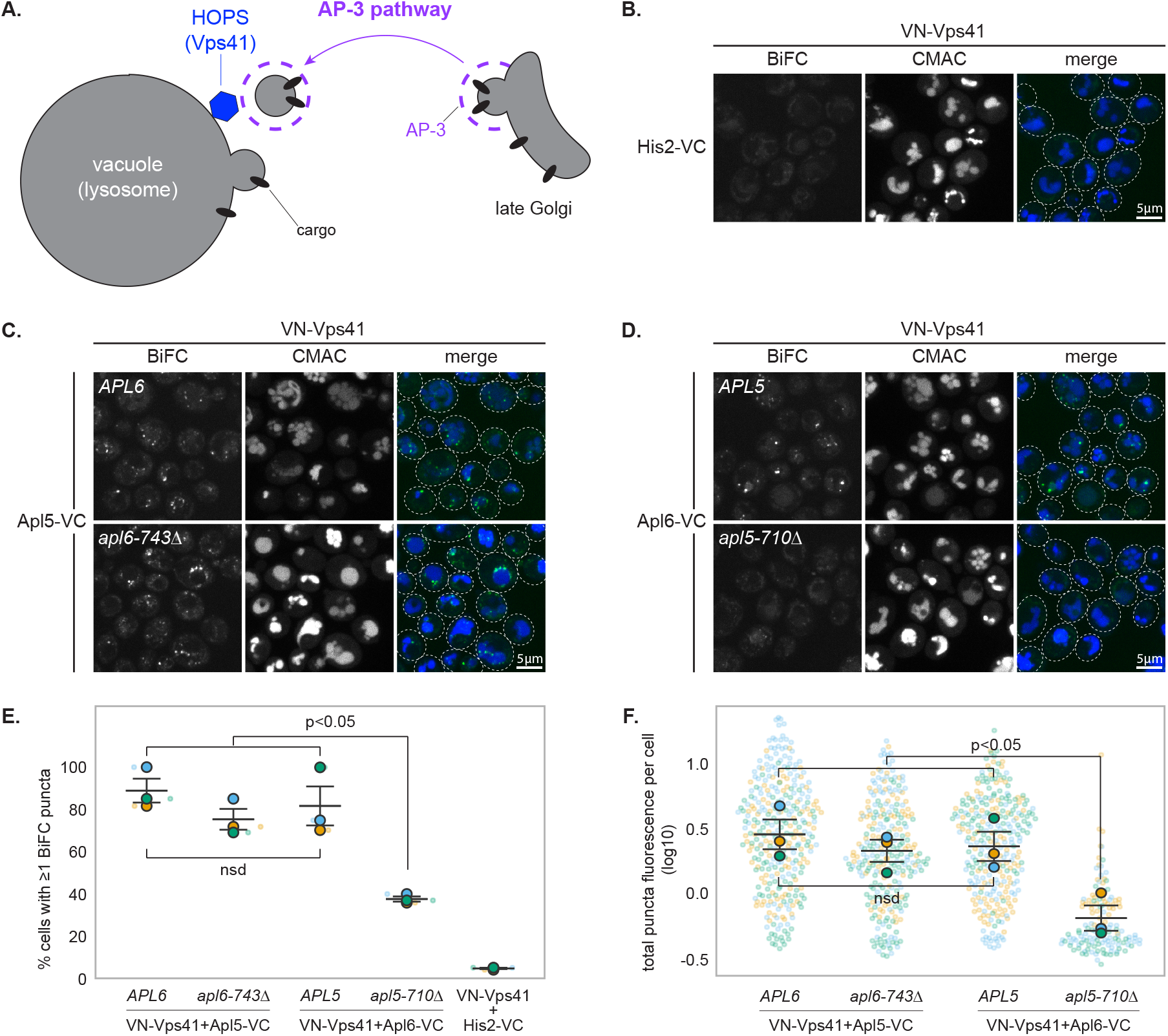
The Apl5 δ subunit IDR promotes Vps41 engagement with AP-3. (A) Schematic showing HOPS complex at the vacuole and its established direct interaction with AP-3. Vps41 of the HOPS complex binds directly to the Apl5 IDR (Angers and Merz, 2009; Schoppe et al., 2020). (B-D) Representative confocal microscopy images of BiFC between VN-Vps41 (expressed from centromeric plasmid) and the indicated genomically-integrated VC fusions in wild-type or IDR deletion (*apl5-710Δ* or *apl6-743Δ*) backgrounds. His2-VC (B) serves as negative control. VN-Vps41 produced no signal with His2-VC but robust BiFC puncta with both Apl5-VC and Apl6-VC in wild-type cells, confirming the known Vps41-Apl5 interaction. While Vps41-Apl5 BiFC was expected based on direct binding, the Vps41-Apl6 signal likely reflects indirect proximity mediated through AP-3 spatial organization (Figure 1) rather than direct binding. Dashed ovals indicate cellular outlines. Scale bars, 5 μm. (E) Percentage of cells containing at least one BiFC punctum (≥50 cells scored per replicate; quantification as in Figure 1E). The Apl6 IDR deletion (*apl6-743Δ*) did not affect VN-Vps41/Apl5-VC association, whereas Apl5 IDR deletion (*apl5-710Δ*) reduced VN-Vps41/Apl6-VC association (puncta-positive cells: 81% to 37%, p<0.05), demonstrating that the Apl5 δ subunit IDR serves as the primary organizer of the AP-3-HOPS interface. AP-3 spatial organization potentially amplifies this interaction by bringing Apl6 into proximity with Vps41-bound Apl5. (F) Total puncta fluorescence per puncta-positive cell (quantification as in Figure 1F).

**Supplemental Figure S3.**
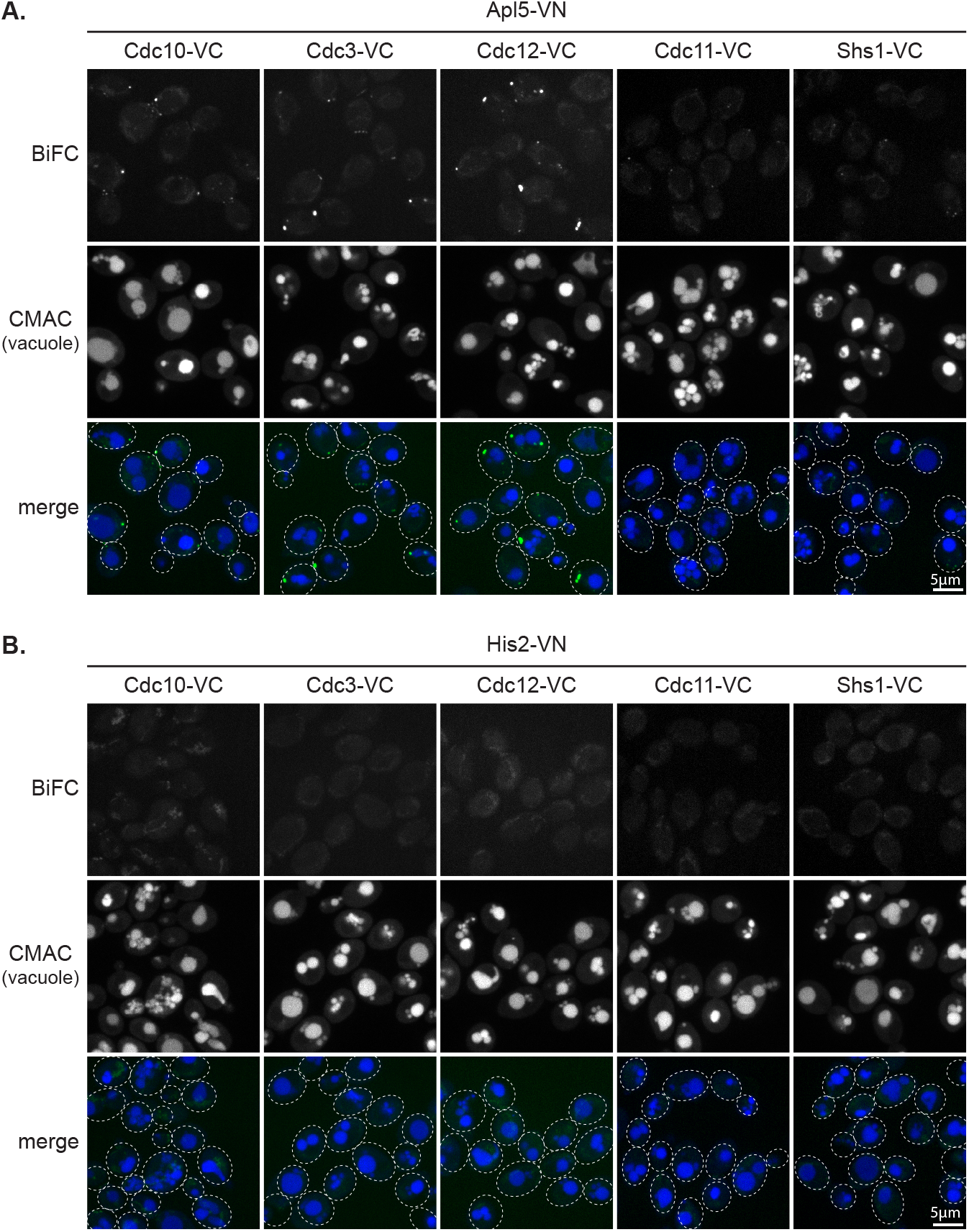
AP-3 shows hierarchical BiFC proximity to septin subunits. (A) Representative confocal microscopy images of Apl5-VN paired with septin-VC fusions (Cdc10-VC, Cdc3-VC, Cdc12-VC, Cdc11-VC, Shs1-VC). Robust puncta are observed with core septins (Cdc10, Cdc3, Cdc12) while terminal septins (Cdc11, Shs1) show diminished signals, mirroring the hierarchy observed with Apl6-VN in Figure 3B. The differential BiFC pattern reveals that AP-3 shows preferential proximity to core septin subunits that form the central architectural scaffold of septin octamers. Core subunits Cdc10, Cdc3, and Cdc12 form the palindromic center of octamers, whereas Cdc11 and Shs1 occupy terminal positions (see Figure 3A). The consistent hierarchy observed with both Apl5-VN and Apl6-VN pairings (Figure 3B) supports the conclusion that AP-3 engages assembled septin octamers with preferential access to core architectural elements rather than terminal caps. Dashed ovals indicate cellular outlines in the merged images. Scale bars, 5 μm. (B) Negative controls showing no BiFC signal above background when septin-VC fusions are paired with His2-VN, confirming specificity of AP-3-septin proximity signals. Dashed ovals indicate cellular outlines in the merged images. Scale bars, 5 μm. Representative images from n=3 independent experiments.

**Supplemental Figure S4.**
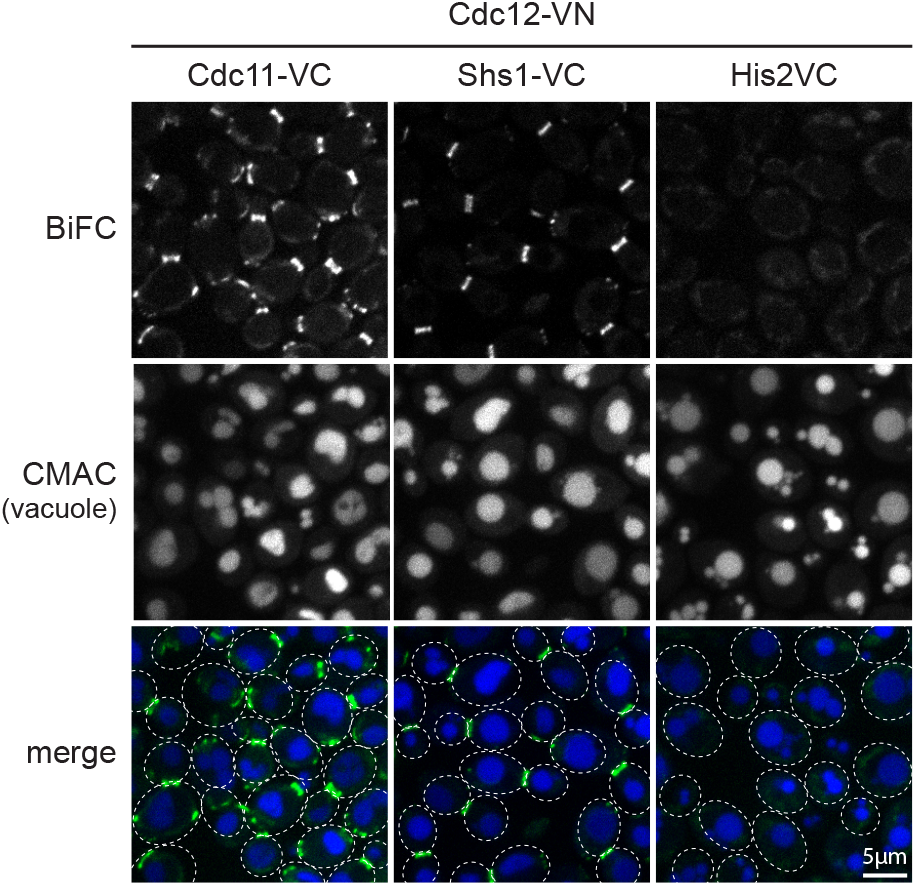
Terminal septin VC fusions are properly expressed and functional. Representative confocal microscopy images showing robust BiFC signal between Cdc12-VN and either Cdc11-VC or Shs1-VC. Because Cdc11 and Shs1 are direct neighbors of Cdc12 within septin octamers (see Figure 3A; Bertin et al., 2008; Garcia et al., 2011; Weems and McMurray, 2017), this BiFC signal confirms that Cdc11-VC and Shs1-VC fusion proteins are properly expressed, correctly folded, and capable of assembling into septin octamers. The terminal septin VC fusions are also capable of participating in productive BiFC reconstitution when their interaction partners are appropriately positioned within ∼7 nm. The robust BiFC observed in these positive control pairings demonstrates that the reduced BiFC signals observed between AP-3 subunits and terminal septins Cdc11-VC or Shs1-VC (Figure 3B-D) reflect genuine differences in spatial proximity or accessibility rather than technical artifacts such as impaired fusion protein expression, misfolding, or inability to reconstitute Venus fluorescence. This control is critical for interpreting the hierarchical interaction pattern, as it establishes that terminal septins are competent for BiFC but simply do not come into close proximity with AP-3 subunits as frequently as core septins do. Dashed ovals inidicate cellular outlines in megred images. Scale bars, 5 μm. Representative images from n=3 independent experiments.

